# TweetyNet: A neural network that enables high-throughput, automated annotation of birdsong

**DOI:** 10.1101/2020.08.28.272088

**Authors:** Yarden Cohen, David Nicholson, Alexa Sanchioni, Emily K. Mallaber, Viktoriya Skidanova, Timothy J. Gardner

**Author notes:** These authors contributed equally to this work (Yarden Cohen, David Nicholson).

## Abstract

Songbirds have long been studied as a model system of sensory-motor learning. Many analyses of birdsong require time-consuming manual annotation of the individual elements of song, known as syllables or notes. Here we describe the first automated algorithm for birdsong annotation that is applicable to complex song such as canary song. We developed a neural network architecture, “TweetyNet”, that is trained with a small amount of hand-labeled data using supervised learning methods. We first show TweetyNet achieves significantly lower error on Bengalese finch song than a similar method, using less training data, and maintains low error rates across days. Applied to canary song, TweetyNet achieves fully automated annotation of canary song, accurately capturing the complex statistical structure previously discovered in a manually annotated dataset. We conclude that TweetyNet will make it possible to ask a wide range of new questions focused on complex songs where manual annotation was impractical.

## Introduction

Songbirds provide an excellent model system for investigating sensorimotor learning [1]. Like many motor skills, birdsong consists of highly stereotyped gestures executed in a sequence [2]. In this and many other ways, birdsong resembles speech: song is learned by juveniles from a tutor, like babies learning to talk [3]. A key advantage of songbirds as a model system for studying vocal learning is that birds sing spontaneously, often producing hundreds or thousands of song bouts a day. This provides a detailed readout of how song is acquired during development, and how this skilled behavior is maintained in adulthood. Leveraging the amount of data that songbirds produce requires methods for high-throughput automated analyses. For example, automated methods for measuring similarity of juvenile and tutor song across development [4,5] led to important advances in understanding the behavioral [6, 7] and genetic [8] bases of how vocalizations are learned. These examples demonstrate how automated methods that enable analysis of large-scale behavioral datasets contribute to realizing the potential of songbirds as a model system.

However, this potential to address central questions of sensorimotor learning is currently hindered by a lack of high-throughput automated methods for scaling up other types of analyses. The central issue is that many analyses require researchers to annotate song. Annotation is a time-consuming process done by hand (typically with GUI-based applications, e.g., Praat, Audacity, Chipper [9–11]). An example of Bengalese finch song annotated with a GUI is shown in Fig. 1. Researchers annotate song by dividing it up into segments (red lines in Fig. 1), often referred to as syllables or notes, and assigning labels to those segments (letters in Fig. 1). Annotation makes several types of analyses possible. For example, annotation is required to build statistical models of syntax [12–15], to fit computational models of motor learning that precisely quantify how single syllables change over the course of an experiment [16,17], and to relate behavior to neural activity [18–20]. Annotating song greatly increases our ability to leverage songbirds as a model system when answering questions about how the brain produces syntax observed in sequenced motor skills, and how the brain learns to adaptively control muscles.

**Fig 1.**
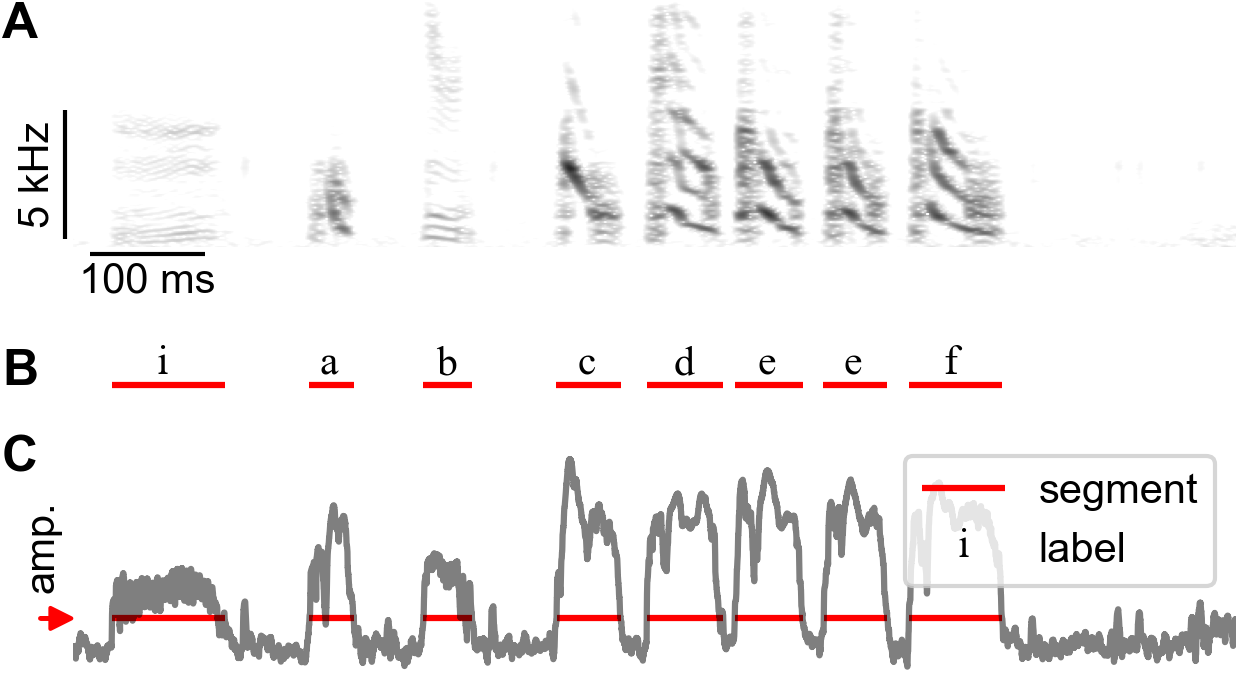
Annotation of birdsong. **A.** Spectrogram showing a brief clip of Bengalese finch song with different syllable types. **B.** Text labels over red segments are applied by human annotators to assign those segments to various syllable classes. **C.** Segments were extracted from song by finding continuous periods above a fixed amplitude threshold. Red arrow to left of panel **C** indicates the user-defined amplitude threshold.

Previous work has been done on automating annotation, as we briefly review below in Proposed Method and Related Work, but these methods are challenged by the variable song of some species. To illustrate these challenges, Fig 2A-C presents examples of annotated songs from different species. When a species’ song consists of just a few syllables sung repeatedly in a fixed motif, methods based on template matching or other algorithms (see Proposed Method and Related Work below) can be applied. This is true for zebra finches, as can be seen in a song from one individual shown in Fig 2A. However, many species have songs that are more complex than the stereotyped motif of zebra finches. Complex songs can contain a large vocabulary of syllable types arranged in multiple motifs or phrases, with phrases sequenced according to complex transition statistics. For example, Bengalese finch song contains “branch points”, where a given syllable may transition to more than one other class of syllable. An example of a branch point is indicated above the spectrogram in Fig 2B. In addition, Bengalese finch song can contain syllables that repeat, with the number of repeats varying from rendition to rendition. Both branch points and repeats prevent existing algorithms from effectively annotating Bengalese finch song (Fig 2E). Canary song is even more complex (Fig 2C). Some individuals may have as many as 50 unique classes of syllables in their repertoire. Bouts of canary song can last more than a minute instead of a few seconds (Fig 2D). These long songs contain individual syllable types that can be very short, under 10ms, or very long, ranging up to 500ms (Fig 2F). Some syllables are very quiet, and others loud. Because of this extreme range of amplitude, common methods for segmenting audio of song into syllables can fail. Segments are typically defined as points where the smoothed sound envelope or other song-related acoustic features [4] stay above some threshold, indicated by the dashed lines in Fig 3. In the case of canary song, if sound energy or other acoustic features are filtered on timescales short enough to accurately segment the shortest syllables, then the longest syllables will be subdivided. This problem is also commonly encountered when analyzing the variable songs of young zebra finches. Fig 3 illustrates how canary song is difficult to segment in an automated manner. Finally, canary song has a hierarchical structure where syllables occur in trilled repetitions, called phrases, that themselves obey long-range syntax rules [12,21]. Phrases can differ in duration depending on the type of syllable being repeated and similarly inter-syllable silent gaps vary widely in duration (Fig. S1). Because of all this complexity, there are currently no automated methods for accurate annotation of canary song.

**Fig 2.**
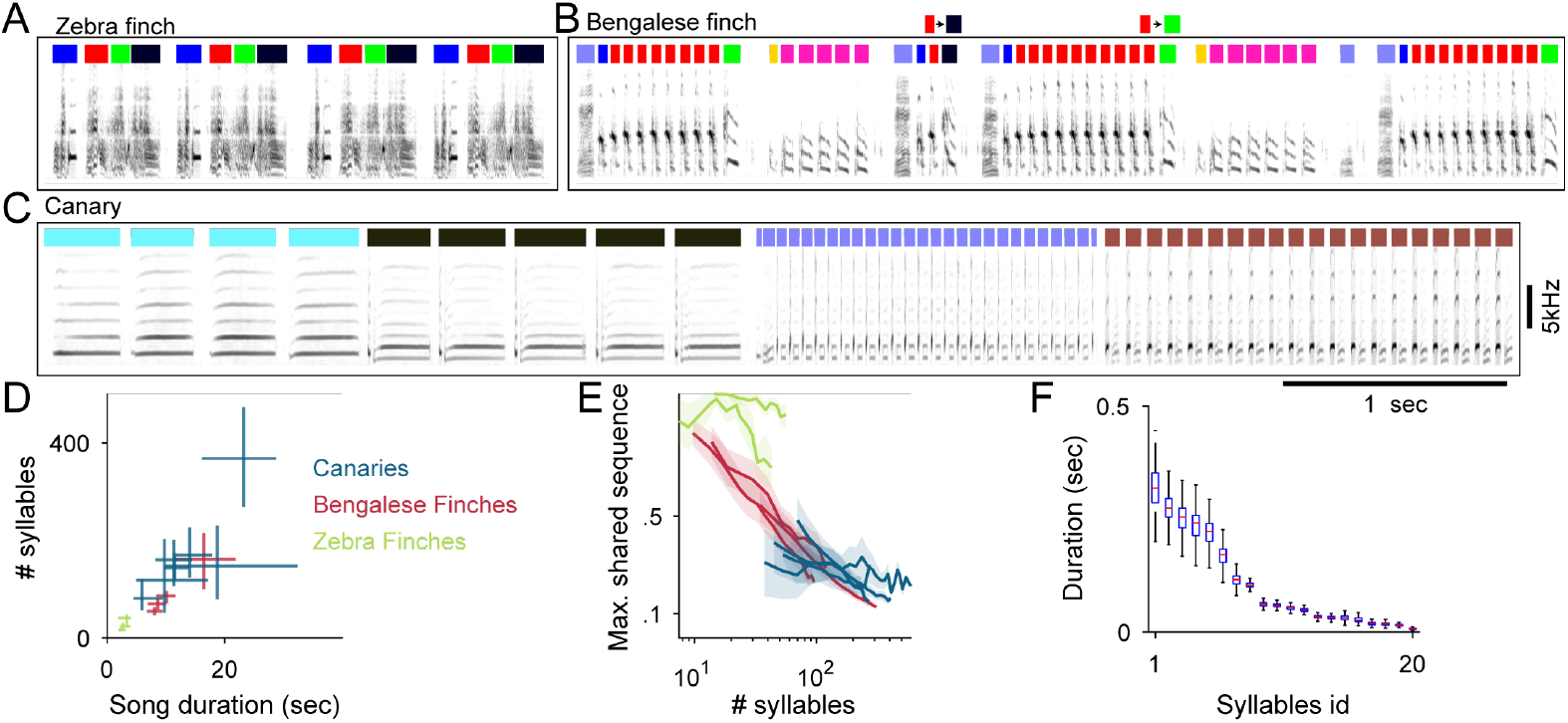
The challenge of annotating complex songs. **A.** The zebra finch repeating motif allows annotation by matching its template spectrogram without segmenting different syllables (colored bars). **B.** Bengalese finch songs segmented to syllables shows variable transitions and changing numbers of syllable repeats. **C.** A third of one domestic canary song of median duration segmented to syllables reveals repetitions (phrase) structure. **D.** The median, 0.25 and 0.75 quantiles of song durations (x-axis) and of syllables per song (y-axis) for 2 canary strains, Bengalese finches and Zebra finches (color coded) **E.** Variable songs are not suited for template matching. Songs contain repeating sequences of syllables but because of sequence variability songs with more syllables (x-axis) share smaller sequence fractions (y-axis) **F.** Distributions of syllable duration for one domestic canary. The bird had 20 different syllable types (x-axis, ordered by mean syllable duration). Box plot shows median, 0.25 and 0.75 quantiles of syllable durations. Whiskers show the entire range.

**Fig 3.**
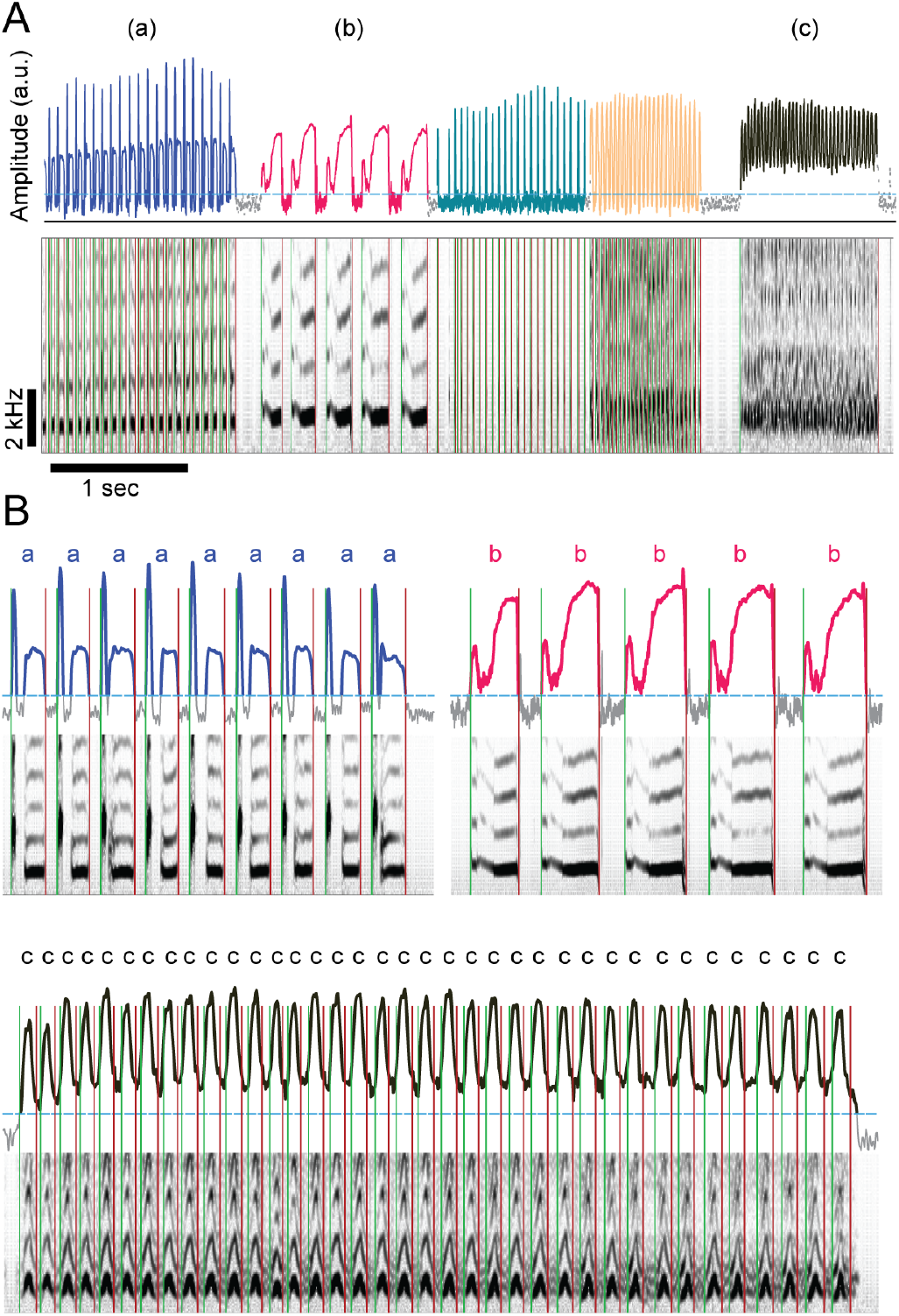
Examples of failure to segment canary song. **A.** Several seconds of domestic canary song, presented as a spectrogram, beneath a plot of a band-pass filtered sound amplitude. To segment song, an amplitude threshold can be taken, marked by the dashed line on the amplitude trace, and then an automated program finds continuous segments of above-threshold amplitude and marks the onset and offset times of those segments (green, red lines in the spectrogram panel). **B.** Focusing on three examples (a-c matching panel A), segmenting by threshold crossing with a fixed filtering bandwidth does not work well for canaries. Above threshold amplitudes are shown in bold colored lines and reveal that syllables of type ‘a’ are broken into 2 components and syllables of type ‘c’ are not separated by low amplitude.

### Proposed Method and Related Work

Previous work has been done to automate annotation, as referenced above, that we now briefly review. The crucial point here is that none of the methods work for canary song, for the reasons we outlined and demonstrated in Figs. 2 and 3, necessitating the development of an algorithm like the one we present. However, for birdsong that consists largely of a single fixed motif, like that of zebra finches, several methods have been widely used, including semi-automatic clustering methods [22,23], and template matching [24–26]. Several studies have also applied supervised learning algorithms to annotation, such as Hidden Markov Models [27], k-Nearest Neighbors [28], and support vector machines [29]. These algorithms can annotate more variable song with branch points and repeats, like that of Bengalese finches, but they all require segmenting song to extract the engineered features used to train the algorithms (e.g. acoustic parameters like pitch and duration). To our knowledge there has been no large-scale comparison of performance of these different algorithms, but at least one study suggests they may not generalize well across songs of different individuals [30]. Additionally, feature extraction can fail if segmentation is noisy, e.g. because of changes in audio equipment set-up, background noises, etc. Here again we stress that canary song exhibits wide ranges in amplitude, and often requires annotators to set multiple thresholds to successfully extract segments. These factors contribute to the lack of automated algorithms for annotating canary song.

Given these issues, we sought to develop an algorithm for automated annotation that (1) can learn features from data, and (2) does not require segmented syllables to predict annotations. To meet both these criteria, we developed an artificial neural network that we call TweetyNet, shown in (Fig 4). TweetyNet takes as input windows from spectrograms of song and produces labels for each time bin of that spectrogram window. TweetyNet requires no pre-processing of song spectrograms - most importantly, segmentation of song into syllables is not needed. Silent gaps between syllables are labelled in training data, and these silent labels are assigned to gaps between syllables when TweetyNet inference is applied to a new song.

**Fig 4.**
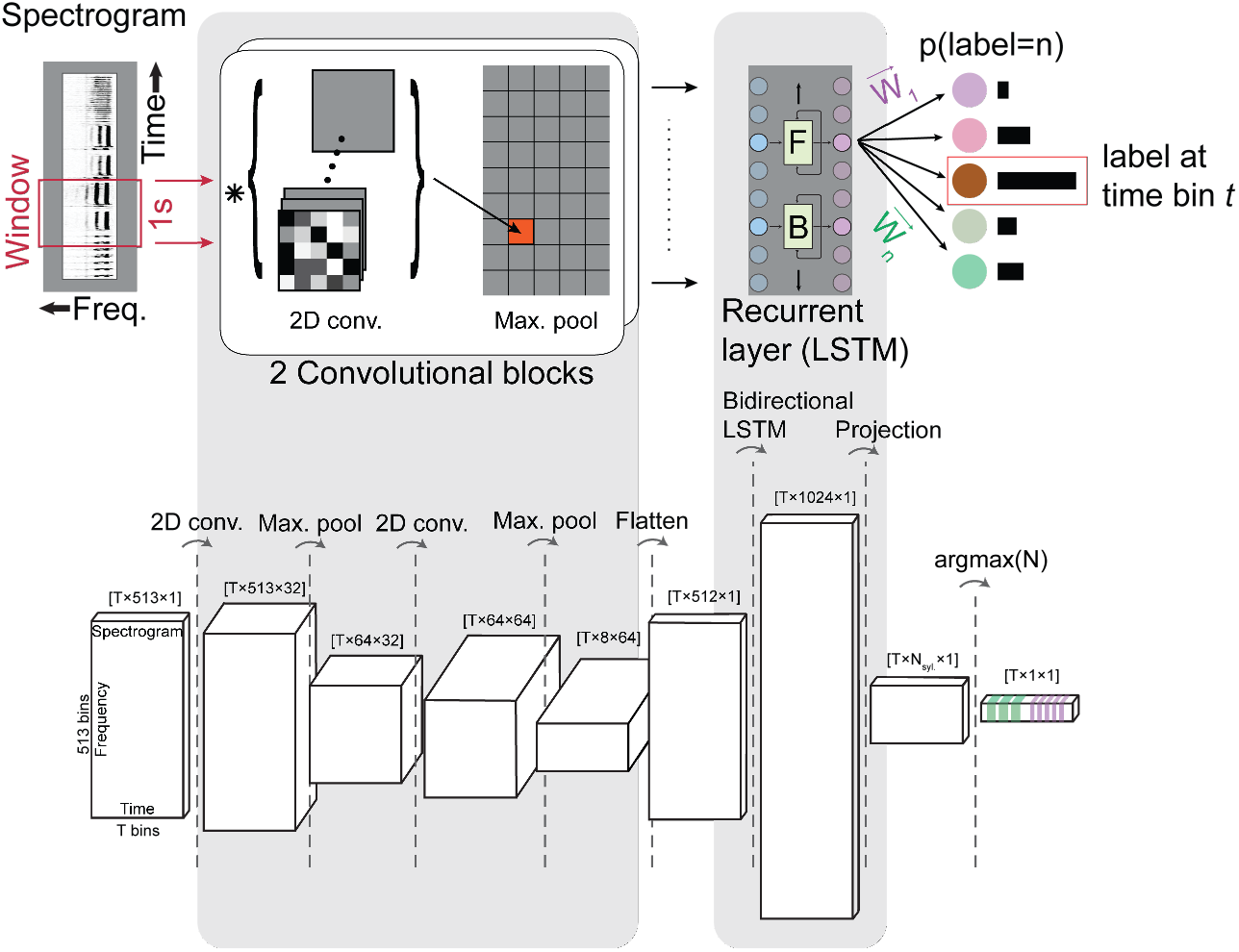
TweetyNet neural network architecture. Top, network schematic. TweetyNet takes as input a window, specified in time bins, from a spectrogram (red box, left) and in a sequence of steps (left to right) outputs a label for each time bin within the window: (1) The convolutional blocks produce a set of feature maps by performing a cross-correlation-like operation (asterisk) between their input and a set of learned filters (greyscale boxes). A max-pooling operation down samples the feature maps. (2) The recurrent layer is made up of Long Short Term Memory (LSTM) units, and the number of units equals the number of time bins in the spectrogram window. This step is designed to capture dependencies across time using both forward (F) and backward (B) passes through time to learn. (3) A projection 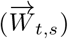 onto the different syllable classes, *s* = 1..*n*, resulting in a vector of probabilities at each time bin *t* that the label is *n*. The number of classes, *n*, is predetermined by the user and includes a class for no-song time bins. (4) Each time bins is labeled by choosing the class with the highest probability and the labelled time bins are used to separate continuous song segments from no-song segments and to annotate each song-segment with a single label. Bottom, the shapes of tensors (multi-dimensional arrays) that result from each operation the network performs.

Essentially, the network combines two types of layers found in neural networks:(1) convolutional layers, common in computer vision tasks to learn features of images [31–33], and (2) recurrent layers, often used to predict sequences [34]. A recurrent layer is a natural choice because the input image or spectrogram is defined by two axes (time and frequency) with very different correlation structure. Specifically, the temporal dimension of songbird vocalization, like music and environmental noises, contains regularities in multiple time scales that are unrelated to the regularities of the frequency axes. The bidirectional LSTM (Long-Short-Time-Memory) recurrent layer is designed to capture these temporal correlations. [35,36].

To predict annotation, we feed consecutive windows from spectrograms to trained networks and then concatenate the output vectors of labeled timebins. Finally, we simply find uninterrupted runs of a single syllable label to annotate song syllables from this framewise classification. As discussed below, this final step can include a “debounce” step that requires a minimum syllable duration and choosing a single label for consecutive time bins not labeled as silence by majority vote. In the rest of the results below we show that this simple method trained end-to-end provides robust predictions of segment onsets, offsets, and labels.

Surprisingly, beyond the work previously cited, we find little research that addresses the problem of learning to classify each time bin of a vocalization, either for human speech or birdsong. The architecture we present here is somewhat similar to early deep networks models for speech recognition, but a crucial difference is that state-of-the-art models in that area map directly from sequences of acoustic features to sequences of words [37]. The success of these state-of-the-art models is attributed to the fact that they learn this mapping from speech to text, **avoiding** the intermediate step of classifying each frame of audio, as has previously been shown [34]. In other words, they avoid the problem of classifying every frame that we set out to solve. The architecture that we develop is most directly related to those that have been used for event detection in audio and video [35,36] and for phoneme classification and sequence labeling [34,38]. The closest prior model for segmenting and labeling birdsong is [39]. Several aspects of that study provide context for the contributions of our work. The authors compared different pipelines that combine a neural network for recognizing syllable segments with Hidden Markov Models that learns to predict syllable sequences, and in this way improve the output of the network. They measured performance of these pipelines on a large dataset of hand-annotated Bengalese finch song which they made publicly available [40].

In summary, the key prior art is the important work of Koumura and Okanoya [39]. This work anticipates the overall structure of our model, but through the integration of multiple distinct components that are individually optimized. In contrast, TweetyNet is a single neural network trained end-to-end, meaning it does not require optimizing multiple models. Below we show that TweetyNet meets our criteria for an algorithm that learns features from data and does not require segmented song to make predictions. To do so we we benchmark TweetyNet on Bengalese finch and canary song, and where possible compare the performance to [39]. Additionally we show that we achieve robust performance: across songs of individuals, which can vary widely even within a species; across many bouts of song from one individual, e.g. across days of song, and; across multiple species. Lastly we show that this performance required only a small amount of manually annotated data to train TweetyNet models accurately enough to recreate and add details to the deep structure of canary syntax.

## Results

### TweetyNet annotates Bengalese finch song with low error rates across individuals

We first set out to test whether our network robustly annotates syllables across a large number of individual birds. To do so, we made use of the publicly available repository of Bengalese Finch song [40], used to benchmark hybrid neural network-HMM models from [39] as referenced in Proposed Method and Related Work. The repository contains song from 10 individual birds, with hundreds of bouts of hand-annotated song for each bird. Each individual’s song had different number of syllables and obeyed a different syntax. To benchmark TweetyNet models on this dataset, we generated learning curves that plot error of the model as a function of the size of the training set (duration in seconds). The learning curves give us an estimate of the smallest amount of hand-labeled training data we would need to obtain the lowest error that the TweetyNet model can achieve. For each bird we split the data into fixed training and test sets, with durations of 900 and 400 seconds respectively. Then for each training set duration we trained 10 replicates with a randomly-drawn subset of the training data. We computed error metrics for each training replicate on the held-out test set for each individual. (See Materials and methods for details.) As shown in Fig 5, these learning curves demonstrate that TweetyNet models achieved low error rates across all ten birds. We first looked at frame error, a percentage that measures the number of times the label predicted by the model for each time bin in a spectrogram did not match the ground truth label. For all birds TweetyNet models achieve less than 8% frame error with the smallest training set duration of 30 seconds (Fig 5A). From the learning curve we can estimate that across birds, the lowest frame error that TweetyNet models produce is roughly 4%, and that they achieve this with just 180 seconds (three minutes) of training data. (For specific values, see Table 1.) Larger training sets did not further reduce error.

**Fig 5.**
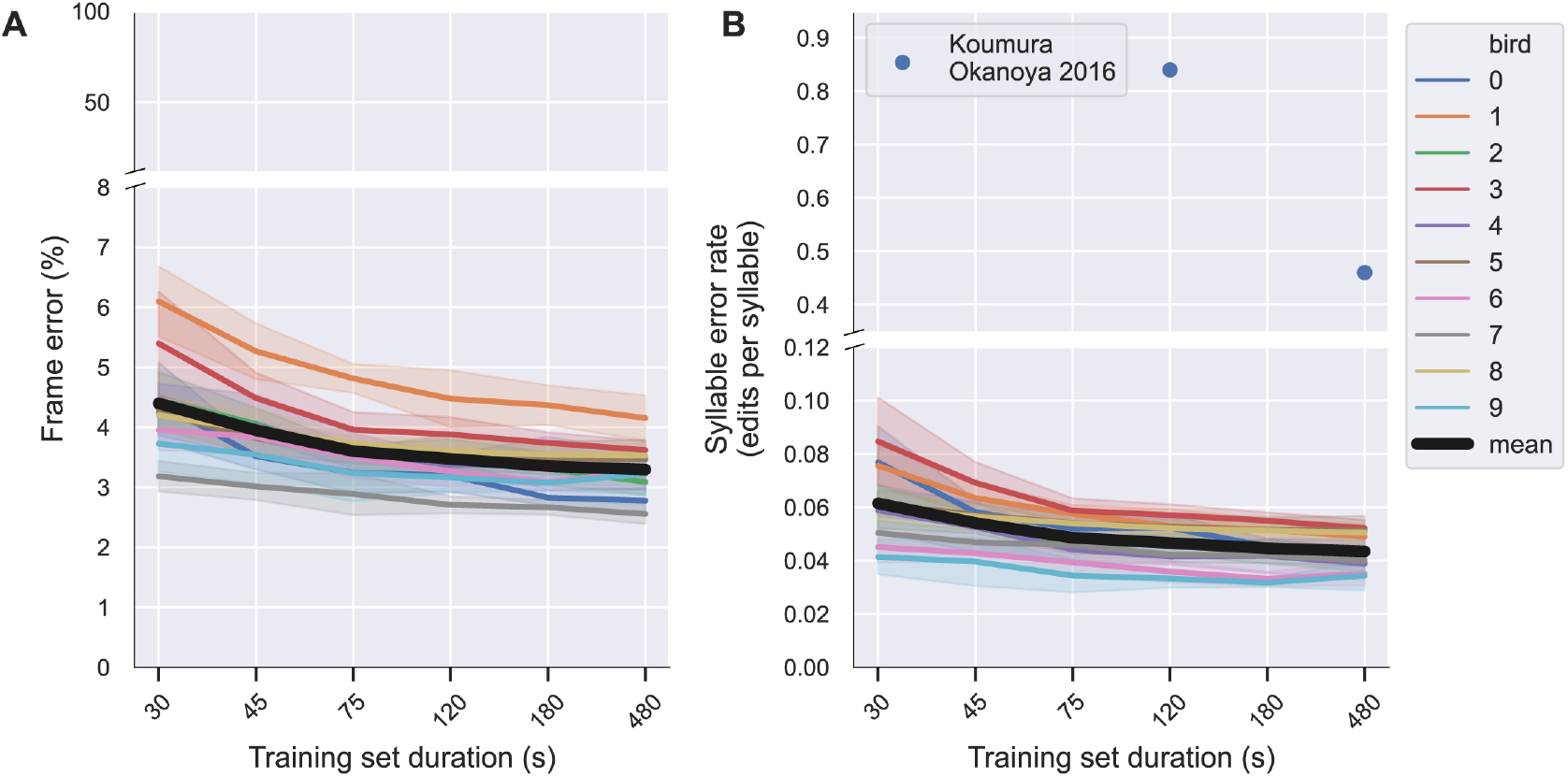
TweetyNet annotates song with low error rates across ten indiviudal Bengalese finches. Model ‘learning curves’ showing the reduction in annotation error (y-axis) on a held-out test set as a function of the size of the training set (x-axis). Shown are the ‘frame error rate’ (**A**) measuring the percent of mislabeled time bins and the ‘syllable error rate’ (**B**) measuring the normalized sequence edit distance. Each colored line corresponds to one bird from dataset. The solid line indicates mean error across ten training replicates for each training set duration, and the translucent error band around the solid lines indicates standard deviation. Thicker black lines indicate the mean across birds. Circular blue markers indicate mean syllable error rate across birds reported in [39] for a different algorithm using the same dataset.

**Table 1.** Error metrics of TweetyNet models for different species and training set sizes. For each Bengalese finch (BF) and canary (Can) data set we evaluate test-set errors metrics for models trained on several training-sets sizes (measured in seconds). Presented are the mean ± standard deviation across all birds and experiment replicates. The *frame error rate* and *syllable error rate* columns present the raw error shown in learning curves (Figs. 5,7). The *syllable error rate* (*majority vote*) column shows the syllable error rate after applying post-hoc cleaning of annotation, where we assigned a single label to each segment by majority vote and discarded all segments below a set duration (methods). The % *Near Boundary* column shows the percent of frame errors involving silent periods that occur within 0-2 time bins of syllable boundaries (onsets and offsets, see Materials and methods).

| dataset | training set duration (s) | frame error (%) | syllable error rate | syllable error rate (majority vote) | % Near Boundary |
| --- | --- | --- | --- | --- | --- |
| B.F. 1 | 120 | 3.5±0.5 | 0.05±0.01 | n/a | n/a |
| B.F. 1 | 180 | 3.4±0.5 | 0.04±0.01 | n/a | n/a |
| B.F. 1 | 480 | 3.3±0.5 | 0.04±0.01 | n/a | n/a |
| B.F. 2 | 180 | 2.9±1.4 | 0.2±0.09 | 0.06±0.04 | 64.9±14.3 |
| Can. | 240 | 3.9±1.0 | 0.155±0.072 | 0.076±0.037 | 51.1±10.7 |
| Can. | 600 | 3.1±0.8 | 0.09±0.016 | 0.051±0.011 | 58.3±11.3 |
| Can. | 6000 | 2.1±0.8 | 0.069±0.013 | 0.031±0.005 | 68.6±13.9 |

To better understand how well the network segments and labels songs, we used another metric, the syllable error rate, which is analogous to the word error rate that is widely used in the speech recognition literature. This metric is an edit distance that counts the number of edits (insertions and deletions) needed to convert a predicted sequence of syllables into the ground-truth sequence. The error rate is normalized by dividing it by the length of the sequences for comparison across birds (e.g. if one bird sang more syllables per bout than another). Measuring the syllable error rate confirmed that TweetyNet consistently achieved similar error rates across the ten birds, as shown in Fig 5B. Because this metric was also used in [39] (as “note error rate”), we can compare our results directly to theirs. As indicated by blue circles in Fig 5B, the best-performing models in that study achieved syllable error rates of 0.83 and 0.46 with two and eight minutes of training data, respectively. TweetyNet always achieved much lower syllable error rates. Taken together, the results from benchmarking TweetyNet on this dataset indicate that the architecture performs well across the song of many individual birds. In addition, it dramatically outperforms existing models with less training data, and does so while being trained end-to-end without requiring optimizations of multiple steps in a pipeline.

### TweetyNet models achieve low error across days even when trained with just the first three minutes of song recorded

We next sought to benchmark TweetyNet in a scenario similar to long-term behavioral experiments for which we hope to automate annotation. For this purpose we used another publicly-available repository [41] with hand-labeled song from four Bengalese finches. Importantly, the repository contains most or all of the songs sung by each bird for multiple consecutive days, as is typically done during a long-term behavioral experiment, and annotation for all those songs (recall that experimenters usually are able to annotate only a limited number). Here we sought to measure how well TweetyNet models would perform when an experimenter takes the *first* set of songs of some duration *n* and annotates those songs manually before using them to train a network. This stands in contrast to the experiment in Fig. 5, where we trained multiple replicates with random subsets of songs from a larger training set, in order to obtain a better estimate of expected error rates. Of course our goal is to avoid the need for experimenters to label a large dataset by hand and then use it to train multiple replicates with random subsets of that data, just to find the best performing network. If we show that we can achieve comparable error rates with just the first n minutes of song, we can be more confident that TweetyNet models will robustly segment and label hours of song recorded across days.

Using the learning curves in Fig 5 we estimated that three minutes of data was the shortest duration training set we could use to obtain the lowest error rate achieved by models. Thus, we trained single TweetyNet models with the first three minutes of song sung by a bird on one day, and then measured the accuracy of that model using all other songs across multiple days. The test datasets we used to obtain these measures were in almost all cases at least as large as those we used to benchmark models in the learning curves. The mean duration of these test datasets was 1528 seconds (standard deviation of 888.6 seconds, i.e. 25 minutes mean, 14 minutes standard deviation), in contrast to Fig 5 where we measured error with a test set of 400 seconds (6 minutes 40 seconds). Hence this approach gave us multiple estimates of how a single trained model performs on relatively large datasets. TweetyNet models trained in this manner did achieve low frame error (Fig 6A) and low syllable error rates (Fig 6B) across days without exhibiting large fluctuations. The frame error ranged from 2-4% across 3-5 days of song, comparable to those observed when training with a random subset of songs, as in Fig 5. In one case, for one bird, the frame error did increase on the last day, but was still within the low end of the range seen for all birds, and this increase did not appear to translate into an increase in the syllable error rate (Fig 6B and Fig 6C, bird ID or60yw70, red line).

**Fig 6.**
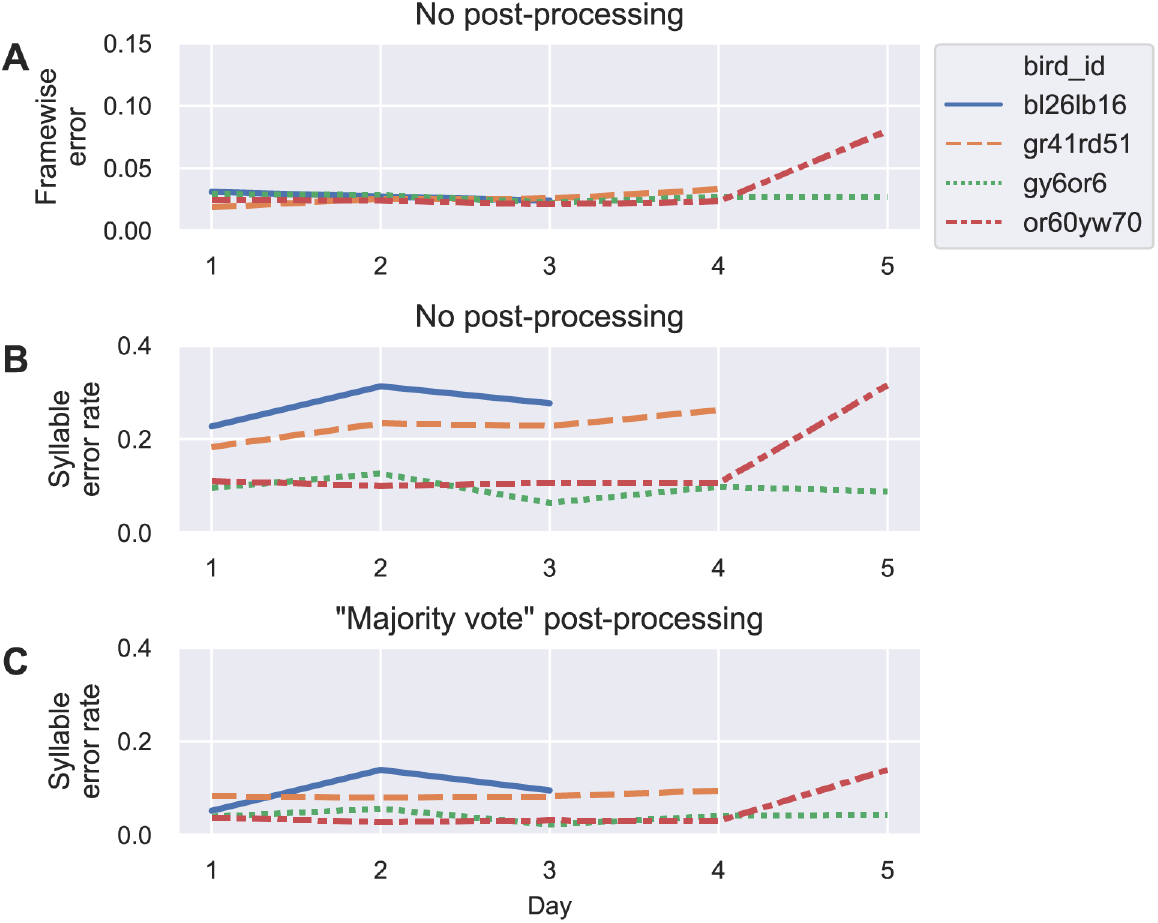
TweetyNet models achieve low error across days of Bengalese finch song, even when trained with just the first three minutes of song recorded. **A.** TweetyNet models trained on the first three minutes of song from day 1 achieved low frame error across days. The mean duration of the set of songs for each day that we used to measure error was 1528 seconds(888.6 S.D.), (i.e. 25 minutes (14 minutes S.D.)), Different line colors and styles indicate individual birds **B.** TweetyNet models trained on the first three minutes of song from day 1 also demonstrate a low syllable error rate across days. **C.** The syllable error rates in **B** further improve after applying a “majority vote” post-processing (assigning the dominant label in each continuous segment of time bins not annotated as ‘silence’, see methods). For one bird (or60yw70), the error did increase on the last day, but was still within the low end of the range seen for all birds.

We also found that TweetyNet models trained on the first three minutes of song maintained a low syllable error rate across days (Fig 6B and Fig 6C), again comparable to what we observed in the learning curves (Fig 5B). Here we additionally tested whether a simple post-processing step could further lower the error rate. This “majority vote” transform consists of taking each labeled segment (bordered by two segments that the network predicted were “unlabeled” / “silent” segments), finding the label occurred most frequently within that segment, and then assigning that label to all time bins within the segment. As shown in Fig 6C, this simple post-processing step did lower the syllable error rate of TweetyNet models. We did not find that this post-processing step had a large effect on the frame error (not shown in plot), from which we infer that this transform removes small frame errors (e.g. a single time bin) that give rise to spurious extra segments, and correcting these in turn produces a large drop in the syllable error rate. Hence we have shown using Bengalese finch song that TweetyNet outperforms existing models and that, with only minimal cleaning of its output, analyses of behavioral experiments can be scaled up to very large datasets.

### TweetyNet annotates minutes-long canary songs with low error rates across individuals

After demonstrating TweetyNet’s high performance across multiple individuals of the same species and across multiple songs of individual birds, we wanted to test TweetyNet across species. We chose the domestic canary (serinus canaria) - a species for which there are no published annotation algorithms and whose rich song repertoire offers a unique opportunity for neuroscience research [12,21,42–44].

As in our first test in Bengalese finches, we curated training sets of 1-10 minutes of song from 3 canaries and measured the frame error rates in a held-out test set 20-30 minutes long. (Training sets are relatively longer than the Bengalese tests since canary songs can be up to a minute or more in length and even sparse sampling of the full repertoire requires these longer training sets.) Still, Fig 7 shows that in three canaries the model learning curves asymptote with 8-10 minute training sets to frame error rates similar to TweetyNet’s performance in Bengalese finches.

**Fig 7.**
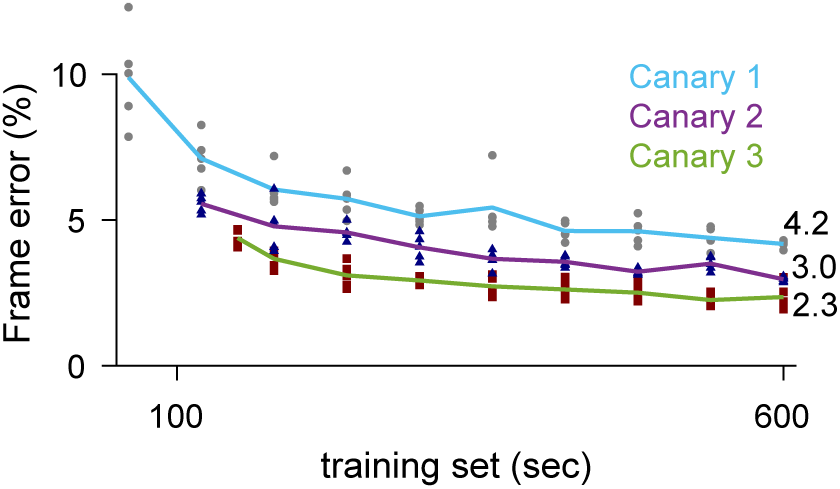
TweetyNet segments and labels canary song with low error rates, similar to Bengalese finches, across individuals. Models were trained on 60s-600s of song from each individual. The mean frame error (lines) of five models (markers) trained with different randomly-drawn subsets from the training set was measured on a separate 1500-2000s test set from each individual. The asymptotic error rates, annotated to the right of the curves, overlaps with the error rates in the Bengalese finch data sets

Unlike TweetyNet’s performance in Bengalese finches, the frame error rates in annotating canary songs cannot be compared to alternative algorithms using published data and results. Furthermore, the length of these songs, usually containing hundreds of syllables, mean that even in very low error rates we expect annotation errors in many songs (Table 1, Fig. S2). These annotation errors can occur at the onset of song and in transitions between canary phrases (Fig 8) and affect analyses of canary syntax.

**Fig 8.**
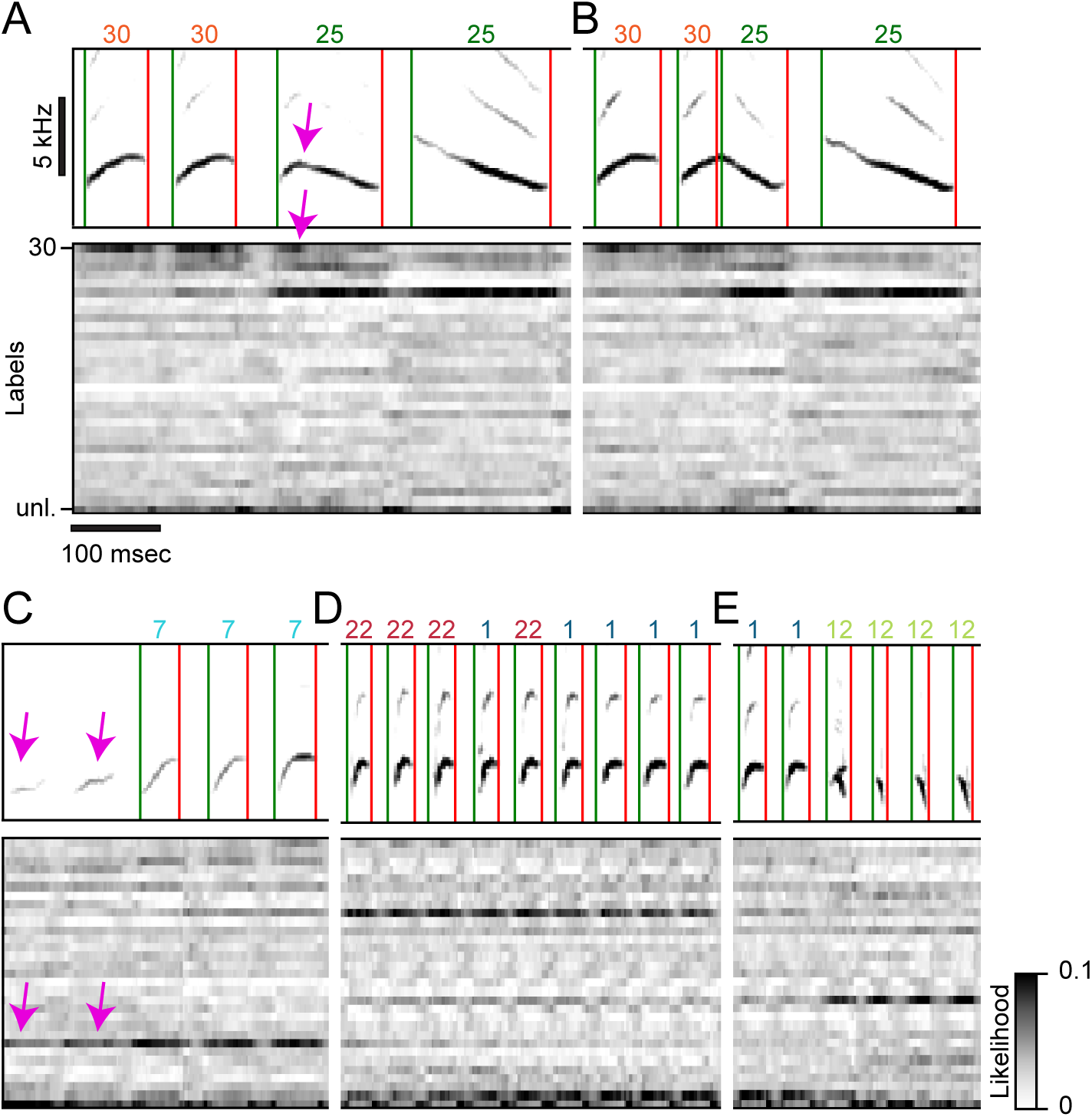
Variants of canary song introduce segmentation and annotation errors. Canary vocalizations contain variations that challenge TweetyNet. The examples in panels A-E show spectrograms on top of the time-aligned likelihood (gray scale) assigned by a trained TweetyNet model to each of the labels (y-axis, 30 syllable types and the tag *unl*. for the unlabeled segments). Green and red vertical lines and numbers on top of the spectrograms mark the onset, offset, and labels predicted by the model. **A,B.** Transitions between syllables can occur without a silence gap. In this example, TweetyNet assigns higher likelihood to both syllables (c.f. pink arrow). In rare variants the model ignores the first syllable (A) **C.** Syllables produced weakly or deformed still get higher likelihood (arrows) but may still be ignored because the unlabeled class gets a higher likelihood. **D.** Transition between phrases of very similar syllables (22 →1) introduce label confusion. **E.** Canaries can produce completely overlapping syllables. The model assigns high likelihood to both classes but is forced to choose only one

To gauge the effect of such errors, in the next section we evaluate the accuracy of syntax models estimated from TweetyNet’s automatic annotation.

### Automated analysis of canary song structure

Sequences of canary phrases contain transitions with different ‘memory’ depths. Namely, the probability distribution of transition outcomes from a given phrase is captured by Markov chains with variable lengths. As shown in a recent study in Waterslager canaries, this syntax structure is captured parsimoniously by probabilistic suffix trees (PST) [12, 45]. The root node in these graphical models, appearing in the middle of Fig 9A,B and Fig 10A,B, represents the zero-order Markov, or base rate, frequencies of the different phrases, labelled in different colors and letters. Each branch, emanating from the colored letters in Figs 9,10 represents the set of Markov chains that end in the specific phrase type designated by that label. For example, the ‘A’ branch in Fig 9a includes the first order Markov model ‘A’ and the second order Markov chains ‘FA’ and ‘1A’ representing the second order dependence of the transition from phrase ‘A’. These models are built by iterative addition of nodes up the branch to represent longer Markov chains, or a transition’s dependence on longer sequences of song history.

**Fig 9.**
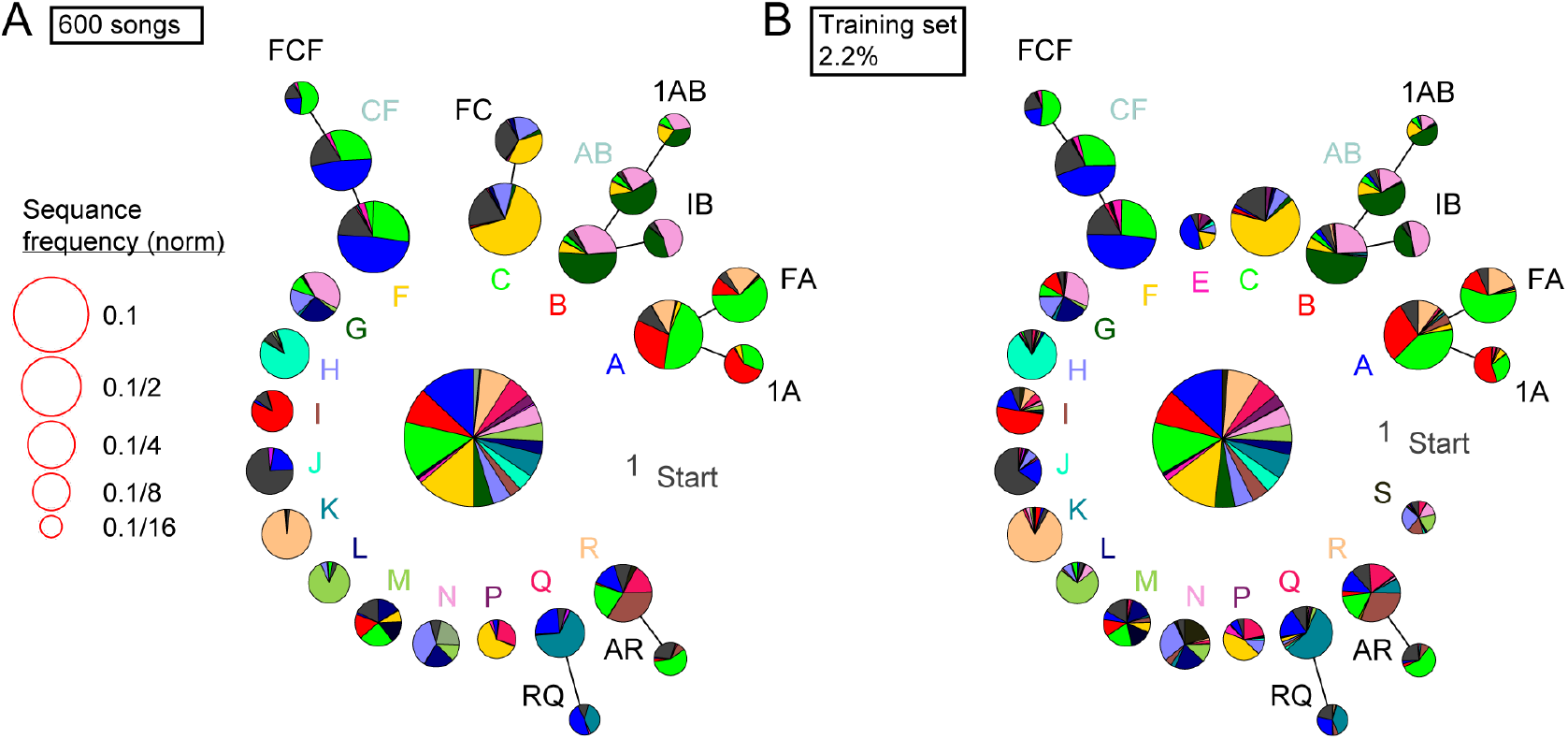
Example of reproducing long-range syntax dependencies, seen in *Waterslager* canaries, in another strain using a TweetyNet model trained on a small fraction of the data. **A.** Long-range order found in 600 domestic canary songs annotated with human proof reader (methods, similar dataset size to [12]). Letters and colors label different phrase types. Each branch terminating in a given phrase type indicates the extent to which song history impacts transition probabilities following that phrase. Each node corresponds to a phrase sequence, annotated in its title, and shows a pie chart representing the outgoing transition probabilities from that sequence. The nodes are scaled according to their frequency (legend). Nodes that can be grouped together (chunked as a sequence) without significantly reducing the power of the model are labeled with blue text. **B.** The songs used to create the PST in A are a subset of 1764 songs. A TweetyNet model was trained using about 2.2% of that dataset (about 9.5% of the data in A). The PST created from the model’s predicted annotation of the entire dataset is very similar to A.

**Fig 10.**
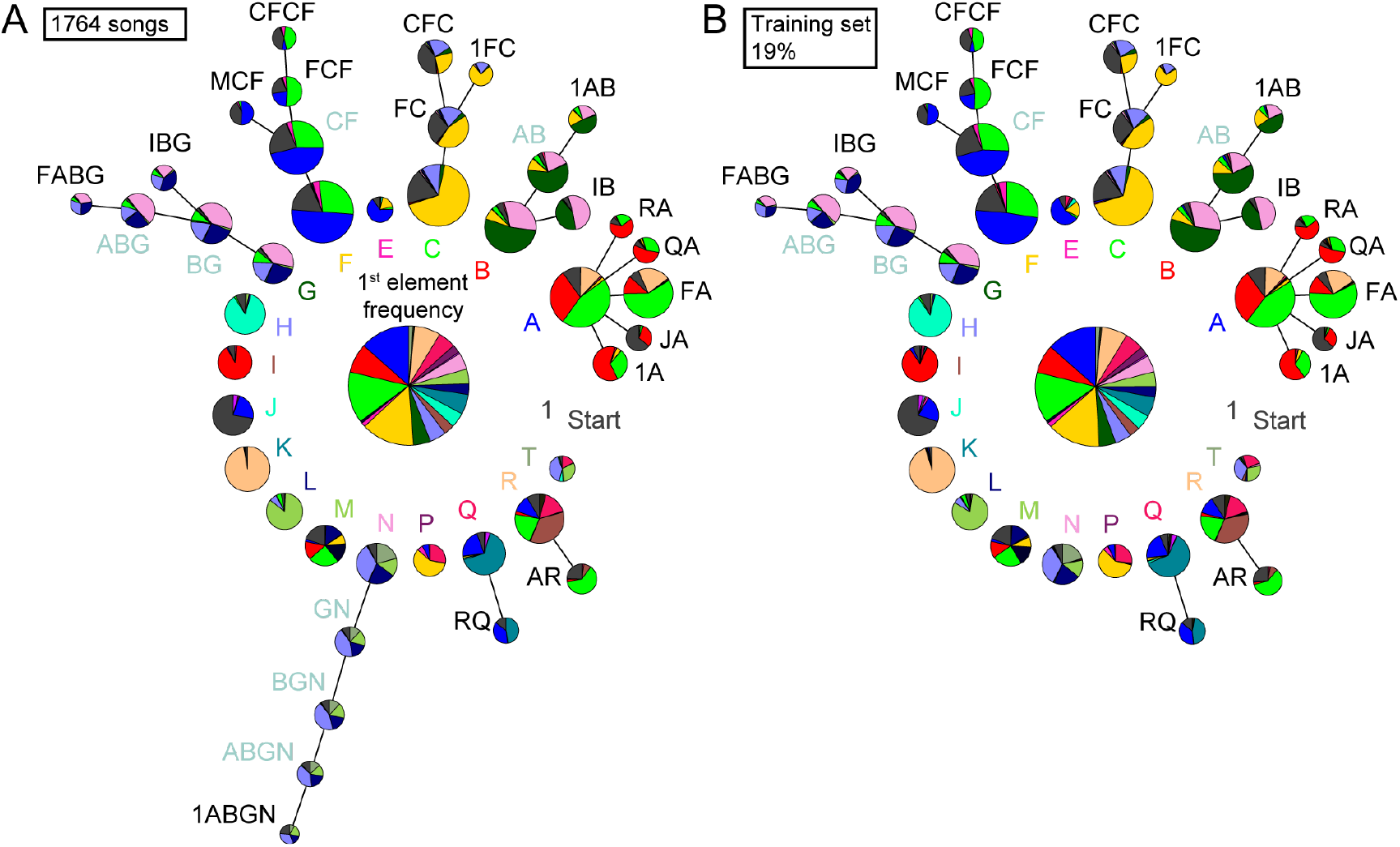
Example of how using TweetyNet to process a larger dataset of canary song adds detail and limits the memory of the syntax structure. **A.** The full dataset of 1764 songs from Fig 9, annotated with a human proof reader, allowed creating a PST with greater detail. Compared to Fig 9A, some branches did not grow. **B.** An almost identical PST was created *without* a human proof reader from a TweetyNet model trained on 19% of the data. The fluctuation in transition probabilities accumulates in long sequences and, in this example, increased the minimal sequence probability included in the PST. This difference prevented the inclusion of the ‘N’ branch.

Figures 9 and 10 demonstrate that TweetyNet parses domestic canary song with an accuracy sufficient to extract its long-range order. In both of these figures, we set parameters of the PST estimation algorithm to derive the deepest syntax structure possible without overfitting as practiced in a recent study [12] that used about 600 hand annotated songs of Waterslager canaries. In this example, using 2.2% of the data set, about 40 songs, to train a TweetyNet model and predict the rest of the data reveals the deep structures shown in Fig 9B - comparable to using 600 hand annotated songs of the same bird (Fig 9A). With more training data, Tweetynet’s accuracy improves as does the statistical strength of the syntax model. In Fig 10B a TweetyNet model was trained on 19% of the data, about 340 songs, and predicted the rest of the data. The resulting syntax model can be elaborated to greater depth without overfitting. To crosscheck this deeper model, we manually annotated all 1764 songs of that bird, revealing a very similar syntax model (Fig 10A).

In sum, we find that TweetyNet, trained on a small sample of canary song, is accurate enough to automatically derive the deep structure that has formed the basis of recent studies [12,46].

### Larger data sets of annotated canary song add details and limit the memory of the syntax structure

The increase in syntax detail, presented in Fig 10, is possible because more rare nodes can be added to the PST without over-fitting the data. Formally, the PST precision increase in larger data sets is defined by the decrease in minimal node frequency allowed in the process of building PST models (Fig 11), as measured in model cross validation (methods). In our data set, we find an almost linear relation between the number of songs and this measure of precision - close to a tenfold precision improvement.

**Fig 11.**
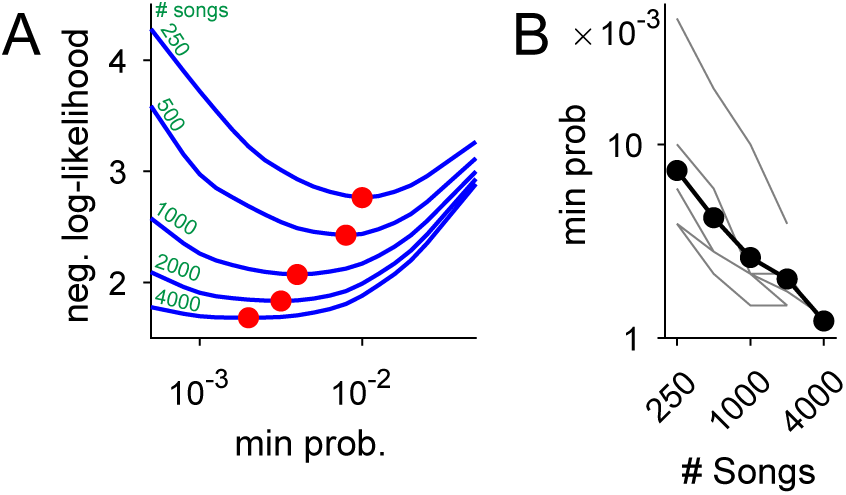
Using datasets more than 5 times larger than previously explored increases statistical power and the precision of syntax models. **A.** Ten-fold cross validation is used in selection of the minimal node probability for the PSTs (x-axis). Lines show the mean negative log-likelihood of test set data estimated by PSTs in 10 repetitions (methods). Curves are calculated for datasets that are sub sampled from about 5000 songs. Red dots show minimal values - the optimum for building the PSTs. **B.** The decrease in optimal minimal node probability (y-axis, red dots in panel A) for increasing dataset sizes (x-axis) is plotted in gray lines for 6 birds. The average across animals is shown in black dots and line.

In Fig 10A, this increased precision allowed reliably adding longer branches to the PST to represent longer Markov chains (in comparison to Fig 9A). In this example, using a dataset 3 times larger revealed a 5-deep branch that initiate with the beginning of song (’1ABGN’) indicating a potential global time-in-song dependency of that transition. The PST in Fig 10A also has branches that did not ‘grow’ when more songs were analyzed (e.g. the ‘B’, ‘Q’, and ‘R’ branches) - indicating a potential cutoff of memory depth that is crucial in studying the neural mechanisms of song sequence generation.

The data sets used in Figs 9A, 10A, and Fig 11, are about 10 times larger than previous studies. To ascertain the accuracy of the syntax models, in creating the data sets we manually proof read TweetyNet’s results (see methods). Across 5 different human proof readers we compare the time required to manually annotate canary song with the proof reading time and find that using TweetyNet saves 95-97.5 percent of the labor.

Taken together, the TweetyNet algorithm allowed us to annotate many more songs of individual complex singers than previously demonstrated, with high accuracy across individuals and across species. This accuracy allowed fully-automated analyses, saved most of the labor, and revealed novel details of canary syntax in a new strain.

## Discussion

The family of songbirds that learns by imitation consists of over 4500 species of birds. Some of these singers, such as the canary, produce songs that are much too complex to be automatically annotated with existing methods, and for these complex singers little is known about the syntax structure and organization of song. Even for birds with simple adult songs, a detailed description of song development will require the application of new methods. This is particularly true for early song development where template based extraction of song syllables and clustering of syllable forms provides an incomplete picture of the full variability of song.

A recent study illustrated the surprises that await a more detailed analysis of song. The canary, one of the most widely bred species of domesticated songbird, was recorded for 2 hours or more and hundreds of songs were manually annotated and cross validated. This data set revealed a new complexity to the statistical structure of canary song - the song follows long-range rules where specific subsets of syllables follow transition statistics governed by 4th and 5th order Markov processes in phrase types [12]. This rich behavior motivated another recent study to implant miniature microscopes in singing canaries, and the recorded neural signals included hierarchical memory traces corresponding to the complex syntax [46]. The sophistication of the neural representation of song in canaries was largely unanticipated based on decades of neural recordings in simpler singers.

The present project was motivated by these recent studies and the knowledge that new fundamental discoveries in vocal learning and neural dynamics will follow if automated annotation of complex song becomes possible. Some methods for automated annotation exist, but previous work suggests these methods have their own limitations, especially when applied to song with many syllable types and variable sequence such as that of Bengalese finches and canaries.

The TweetyNet algorithm described here is a work in progress, with many clear paths for improvement. Still the first syllable error rates described here are dramatic improvements over a prior model for song parsing. We used publicly-available datasets of Bengalese finch song to benchmark TweetyNet. We showed that it achieves low error rates across many individuals. On Bengalese finch data, our single network trained end-to-end performs better than a previously proposed hybrid HMM-neural network model and does so with less training data (Fig. 5). We then showed that TweetyNet models achieve low error across days, in thousands of Bengalese finch songs, even when trained with just the first three minutes of song. This experiment, while strongly restricting the data available for model training, demonstrates the usefulness of TweetyNet in ‘real-life’ laboratory settings - for experimentalists that want to hand annotate as little as possible.

We next reported that TweetyNet was sufficiently accurate to reproduce the recent findings on the complex syntax structure of canary song with fully automated machine-classified song. Specifically, a TweetyNet model trained on just 10 minutes of canary song could accurately recover the statistical structure reported from 600 manually annotated songs - exceeding 100 minutes. Furthermore, a deep network trained on 340 annotated songs, about 19% of the data, could classify a larger data set of more than 1700 songs and build a much more complete statistical model of song revealing additional depth to the long-range syntax rules and extending prior reports on the complexity of the canary song behavior. This more complex statistical model was validated using a manually curated data set of all songs.

With a trained model performing at this level it becomes feasible to examine the effect of social context on song syntax, circadian variations in syntax, or the effects of distinct neural perturbations that could effect song syntax while keeping syllable forms intact. On top of sequence variations, many song studies require syllable similarity metrics to examine the effects of such neural or song pertubations, or the ontogeny of syllable forms through development. Here we used TweetyNet to classify the most likely syllable in every time point, focusing not on variations in syllable form but the sequential structure of song syntax. But, the syllable classification is the final processing step in TweetyNet achieved by maximum a-posteriori (MAP, or argmax) estimation following the calculation of similarity to all possible syllables. Thus, the full likelihood function that TweetyNet produces prior to classification may itself be a useful metric for syllable structure, allowing for example the time course of syllable form to be examined through development or as a result of neural perturbations. A syllable similarity metric that can be assigned at each point in time or frame of a spectrogram without syllable segmentation is, by itself, a new development in the field and can be used, in future development, to improve TweetyNet and to apply it to many more species whose song is difficult to segment.

To make TweetyNet useful to a large research community, we developed the `vak` library - a user-friendly toolbox that enables researchers to apply TweetyNet simply by adapting existing configuration files. This library does not require extensive programming knowledge or expertise in neural networks. The framework will allow users to explore different methods of optimizing neural network models that might improve segmentation, and also generate alternative architectures that could incorporate distinct features and topologies. For example, in many domains transformer networks have recently replaced LSTMs for sequence processing. Substituting transformer layers for the LSTM layer could provide advances here. [47]. Aspects of other deep networks applied to animal motor control may improve TweetyNet. Examples include object detection architectures [48,49] applied to mouse ultrasonic vocalizations and animal motion tracking, and generative architectures applied to birdsong and other vocalizations [50–52]. Lastly we note that in principle TweetyNet and `vak` library can be applied to any other annotated vocalization, including calls of bats, mouse ultrasonic vocalizations, and dolphin communication. We do not claim to have achieved the best possible method for automated annotation of vocalizations with neural networks using supervised learning methods, although we have aimed to establish a strong baseline for the work that will build upon ours. That said, we are confident our method enables songbird researchers to automate annotation required for analyses that address central questions of sensorimotor learning.

## Materials and methods

### Ethics declaration

All procedures were approved by the Institutional Animal Care and Use Committees of Boston University (protocol numbers 14-028 and 14-029). Song data were collected from n = 5 adult male canaries. Canaries were individually housed for the entire duration of the experiment and kept on a light–dark cycle matching the daylight cycle in Boston (42.3601 N). The birds were not used in any other experiments.

### Data availability

Open datasets of annotated Bengalese finch song are available at <https://figshare.com/articles/BirdsongRecognition/3470165> and <https://figshare.com/articles/Bengalese_Finch_song_repository/4805749>. Audio data sets of canary song are available from the corresponding author on request.

### Code availability

The code implementing the TweetyNet architecture, and code to reproduce figures in this paper, are available at <https://github.com/yardencsGitHub/tweetynet> (version 0.4.3, 10.5281/zenodo.3978389). To aid with reproducibility of our experiments, and to make TweetyNet more accessible to researchers studying birdsong and other animal vocalizations, we developed a software library, `vak`, available at <https://github.com/NickleDave/vak>. Both TweetyNet and `vak` are implemented using the following opensource scientific Python libraries: torch [53], torchvision [54], numpy [55,56], scipy [57], dask [58], pandas [59], matplotlib [60,61], seaborn [62], jupyter [63], attrs [64] and tqdm [65].

### Data collection

#### Use of available datasets

Bengalese finch song is from two publicly-available repositories. The first [40] was used for results in 4 and can be found at <https://figshare.com/articles/BirdsongRecognition/3470165>. It accompanied the paper [39]. The second [41] was used for results in Fig 5 can be found at <https://figshare.com/articles/Bengalese_Finch_song_repository/4805749>. Apart from recordings made for this manuscript we used publicly available datasets of Waterslager canary songs [12], Bengalese finch songs [39] and Zebra finch songs [66].

#### Domestic canary song screening

Birds were individually housed in soundproof boxes and recorded for 3-5 days (Audio-Technica AT831B Lavalier Condenser Microphone, M-Audio Octane amplifiers, HDSPe RayDAT sound card and VOS games’ Boom Recorder software on a Mac Pro desktop computer). In-house software was used to detect and save only sound segments that contained vocalizations. These recordings were used to select subjects that are copious singers (≥ 50 songs per day) and produce at least 10 different syllable types.

#### Domestic canary audio recording

All data used in this manuscript was acquired between late April and early May 2018 – a period during which canaries perform their mating season songs. Birds were individually housed in soundproof boxes and recorded for 7-10 days (Audio-Technica AT831B Lavalier Condenser Microphone, M-Audio M-track amplifiers, and VOS games’ Boom Recorder software on a Mac Pro desktop computer). In-house software was used to detect and save only sound segments that contained vocalizations. Separate songs were defined by silence gaps exceeding 1 second.

### Audio processing

#### Segmenting annotated phrases of Waterslager canaries

The dataset of waterslager canaries was available from a previous project in the Gardner lab [12]. These songs were previously segmented into phrases, trilled repetitions of syllables, and not to individual syllables. To include this data in Fig 2 we needed to break annotated phrase segments into syllable segments. In each segmented phrase, we separated vocalization and noise fluctuations between vocalizations by fitting a 2-state hidden Markov model with Gaussian emission functions to the acoustic signal. The suspected syllable segments resulting from this procedure were proofread and manually corrected using a GUI developed in-house (https://github.com/yardencsGitHub/BirdSongBout/tree/master/helpers/GUI).

#### Preparing data sets of domestic canaries

##### Bootstrapping annotation with TweetyNet

In this manuscript we used annotated domestic canary datasets an order of magnitude larger than previously published. To create these datasets we used TweetyNet followed by manual proofreading of its results. This process, described below, allowed ‘bootstrapping’ TweetyNet’s performance.

Song syllables were segmented and annotated in a semi-automatic process:

- A set of 100 songs was manually segmented and annotated using a GUI developed in-house (https://github.com/yardencsGitHub/BirdSongBout/tree/master/helpers/GUI). This set was chosen to include all potential syllable types as well as cage noises.
- The manually labeled set was used to train TweetyNet (https://github.com/yardencsGitHub/tweetynet).
- In both the training phase of TweetyNet and the prediction phase for new annotations, data is fed to TweetyNet in segments of 1 second and TweetyNet’s output is the most likely label for each 2.7msec time bin in the recording.
- The trained algorithm annotated the rest of the data and its results were manually verified and corrected.

##### Assuring the identity and separation of syllable classes

The manual steps in the pipeline described above can still miss rare syllable types of mislabel syllables into the wrong classes. To make sure that the syllable classes are well separated all the spectrograms of every instance of every syllable, as segmented in the previous section, were zero-padded to the same duration. An outlier detection algorithm (IsolationForest: <https://scikit-learn.org/stable/modules/generated/sklearn.ensemble.IsolationForest.html>) was used to flag and re-check potential mislabeled syllables or previously unidentified syllable classes.

##### Preparing spectrograms inputs for TweetyNet

Spectrograms were created from audio files using custom Numpy (Bengalese finch) or Matlab (canary) code. All spectrograms for song from a given species were created with the same parameters (e.g., number of samples in the window for the Fast Fourier Transform). From initial studies we found that it was necessary to perform standard transforms on spectrograms such as a log transform in order for the neural network to learn. We did not notice any difference in the nature of the transform (i.e, we also used log + 1) although here we do not study this systematically.

### Network Architecture

The network takes a 2D window from a spectrogram as input (red box, left in Fig 4) and produces as output labels for each time bin in the window. The spectrogram window passes through two standard convolutional blocks, each of which consists of a convolutional layer and a max pooling layer. The convolutional layer performs a cross-correlation like operation (asterisk in Fig 4) between the spectrogram window and learned filters (greyscale boxes in Fig 4) to produces feature maps. The max pooling layer uses a similar operation to further reduce feature maps to maximum values within a sliding window (orange bin in Fig 4). Importantly, the window size we use in the max pooling layer has a “width” of one time bin, so that this layer does not down-sample along the time axis (although the convolutional layer does). The output of the second convolutional block passes through a recurrent layer made up of LSTM units, where the number of units equals the number of time bins in the spectrogram window.

The final layer in TweetyNet is a projection 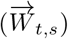 of the recurrent layer’s output onto the different syllable classes, *s* = 1..*n*, resulting in a vector of *n* syllable-similarity scores for each spectrogram time bin *t*. The number of classes, *n*, is predetermined by the user and includes a class for no-song time bins. At present this non-song class includes both background noises and silence, and future iterations of the model may distinguish between these for better performance. To segment syllables, the bin-wise syllable-similarity scores are first used to select a single syllable class per time bin by choosing the label with the highest syllable-similarity score. Since similarity scores can be normalized, this is akin to maximum a-posteriori (MAP) label selection. Then, the labelled time bins are used to separate continuous song segments from no-song segments and to annotate each song-segment with a single label using majority decision.

### Training and benchmarking TweetyNet

Benchmarking of TweetyNet was performed with the `vak` library. We apply standard methods for benchmarking supervised machine learning algorithms, following best practices [67]. We leverage functionality of the `vak` library that extends best practices for benchmarking to the domain where where dataset size is measured in duration, as described in Learning curves.

#### Data transformations

As stated above, the input to the network consists of spectrogram windows. To produce this input, we slid a window of fixed length across spectrograms, essentially creating an array of every possible window from each spectrogram. This array was randomly permuted then fed to the network in minibatches during training, along with the expected output, vectors of labels for each timebin in the spectrogram windows. These vectors of labeled timebins are produced programmatically by `vak` from annotations consisting of segment labels and their onset and offset times. For Bengalese finch song we used windows of 88 time bins, and for canary song we used windows of 370 time bins. We carried out preliminary experiments where we varied the window size for Bengalese finch song, but did not find that larger windows greatly increased accuracy, although they did increase training time.

#### Learning curves

For the studies shown in Figs. 5, 7, we created learning curves, that display a metric such as frame error rate as a function of the amount of training data. For each individual bird we fit networks with training sets of increasing size (duration in seconds) and then measured performance on a separate test set.

In the case of Bengalese finches, we used training sets with durations ranging from 30-480 seconds. For each network trained, audio files were drawn at random from a fixed-size total training set of 900 seconds until the target size (e.g. 60 seconds) was reached. If the total duration of the randomly drawn audio files extended beyond the target duration, they were clipped at the target duration in a way that ensured all syllable classes were still present in the training set. For each bird we trained ten replicates, where each replicate had a different subset of randomly-drawn audio files to create the target training set size. For all Bengalese finches, we measured accuracy on a separate test set with a fixed size of 400s. We chose to use a totally-separate fixed-size set (instead of e.g. using the remainder of the training data set) so we could be sure that any variance in our measures across training replicates could be attributed to the randomly-drawn training set, and not to changes in the test set. We computed metrics such as frame error rate and syllable error rate on the held-out test set for each bird.

For canaries we used test set duration of 1500-2000 seconds and training sets of 60-600 seconds for the learning curves in Fig. 7. For the result in Table 1 we used a test set of 5000 seconds and a training set of 6000 seconds. The method for generating learning curves as just described is built into the `vak` library and can be reproduced using its *learncurve* functionality in combination with the configuration files we shared (reference link) and the publicly-available datasets.

#### Metrics

We measured performance with two metrics. The first is the frame error rate, that simply measures for each acoustic frame (in our case each time bin in a spectrogram) whether the predicted label matches the ground truth label. Hence the range of the frame error rate is between 0 and 1, i.e. can be stated as a percent, and gives an intuitive measure of a model’s overall performance. Previous work on supervised sequence labeling, including bidirectional-LSTM architectures similar to ours, has used this metric [34,38].

The second metric we used is commonly called the word error rate in the speech recognition literature, and here we call it the syllable error rate. This metric is an edit distance that counts the number of edits (insertions and deletions) needed to convert a predicted sequence into the ground-truth sequence. The error rate is normalized by dividing it by the length of the sequences.

In Table 1 we provide two additional measures. The first is a lower bound on the percent of all frame errors that can be attributed to slightly-misaligned syllable onsets and offsets. These syllable boundaries are naturally variable in creating the ground truth hand annotated data sets. Spectrogram time bins in which a trained TweetyNet model and the ground truth disagree and only one of them assigns the ‘unlabeled’ tag can potentially be around segment boundaries. In Fig. S2 we show the histogram of distances, in spectrogram bins, of such frame errors from ground truth segment boundaries. The majority is concentrated in 0-2 bins away from the boundaries, amounting the overall percents summarized in Table 1. The second is syllable error rate after applying post-hoc cleaning of annotation. This cleanup is done in two steps: (1) discard all segments shorter than 5msec (using 10 msec adds an insignificant improvement in some birds) and (2)assign a single label to each segment of time bins not labeled as ‘silence’ by majority vote.

#### Model output as syllable likelihoods

In Fig 8 we present model outputs one step prior to assigning the most likely label to each spectrogram time bin. At that stage, one before the *argmax*(*N*) step in Fig 4, the model output for a given time bin *t* is a real-valued affinity 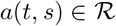 of all predefined syllable classes *s*. In Fig 8 we convert these numbers to likelihoods by subtracting the minimum value and normalizing separately for each time bin 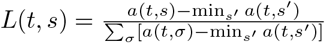. This transformation was done for presentation only. Applying the commonly-used softmax transform 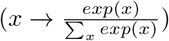 is equivalent since we only keep the maximal value.

### Data analysis - song structure

#### Shared template dependence on number of syllables in song (Fig 2e)

In each bird we define an upper bound for repeating parts of songs using pairwise comparisons. For each song we examined all other songs with equal or larger number of syllables and found the largest shared string of consecutive syllables. The fraction of shared syllables is the ratio between the number of shared sequence and the number of syllables in the first, shorter, song. Then, we bin songs by syllable counts (bin size is 10 syllables) and calculate the mean and standard deviation across all pairwise comparisons.

#### Probabilistic suffix tree (Figs 9, 10)

For each canary phrase type we describe the dependency of the following transition on previous phrases with a probabilistic suffix tree. This method was described in a previous publication from our lab (Markowitz et. al. 2013, code in https://github.com/jmarkow/pst). Briefly, the tree is a directed graph in which each phrase type is a root node representing the first order (Markov) transition probabilities to downstream phrases, including the end of song. The pie chart in Figs 9, 10 shows such probabilities. Upstream nodes represent higher order Markov chains that are added sequentially if they significantly add information about the transition.

### Model cross validation to determine minimal node frequency

To prevent overfitting, nodes in the probabilistic suffix trees are added only if they appear more often than a threshold frequency, *P_min_*. To determine *P_min_* we replicate the procedure in [12] and carry a 10-fold model cross validation procedure. In this procedure the dataset is randomly divided into a training set, containing 90 percent of songs, and a test set, containing 10 percent of songs. A PST is created using the training set and used to calculate the negative log likelihood of the test set. This procedure is repeated 10 times for each value of *P_min_*, the x-axis in Fig 11a. For data sets of different sizes (curves in Fig 11a, x-axis in Fig 11b) the mean negative log-likelihood across the 10 cross validation subsets and across 10 data sets, y-axis in Fig 11a, is then used to find the optimal value of *P_min_* - the minimum negative log-likelihood that corresponds to the highest precision without over-fitting the training set. All PSTs in Figs 9, 10 are created using the cross-validated *P_min_*.

## Acknowledgments

This study was supported by NIH grants R01NS104925, R24NS098536 (T.J.G.) We thank J. Markowitz and T.M. Otchy for sharing song datasets, and Nvidia Corporation for a technology grant (Y.C., Sober lab).

## Supporting information

**Fig. S1.**
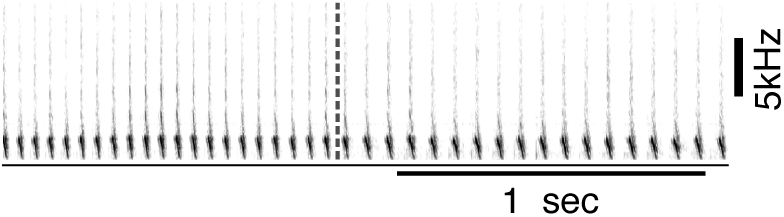
Consecutive canary phrases can include acoustically-similar syllables but differ in the duration of inter-syllabic gaps. Example of two consecutive canary phrases that differ mostly in inter-syllable gaps. In this case, annotation methods that first segment syllables and then use acoustic parameters to classify them will introduce errors. By simultaneously learning acoustic and sequence properties, TweetyNet overcomes this weakness.

**Fig. S2.**
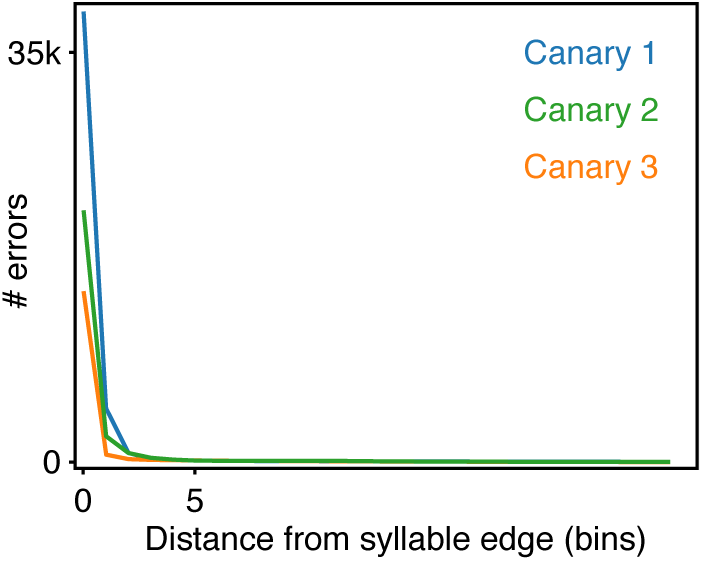
Most errors of trained TweetyNet models are disagreement on syllable boundaries of 0-2 time bins. Potential syllable boundary disagreements are time bins in which the ground truth test set or the trained TweetyNet model disagree and just one of them assigns the ‘unlabeled’ silence tag. The histograms show the distances of those time bins from the nearest syllable boundary in test sets 5000 second long.

## Notes

https://github.com/yardencsGitHub/tweetynet

https://github.com/NickleDave/vak

### Competing Interest Statement

The authors have declared no competing interest.

